# *In vitro* activity of ertapenem against *Neisseria gonorrhoeae* clinical isolates with decreased susceptibility or resistance to extended spectrum cephalosporins in Nanjing, China (2013-2019)

**DOI:** 10.1101/2022.01.25.477800

**Authors:** Xuechun Li, Wenjing Le, Xiangdi Lou, Caroline A. Genco, Peter A. Rice, Xiaohong Su

## Abstract

**Objective:** *Neisseria gonorrhoeae* isolates collected in Nanjing, China, that possessed decreased susceptibility (or resistance) to extended spectrum cephalosporins (ESCs), were examined for susceptibility to ertapenem and their sequence types determined.

**Methods:** Ceftriaxone and cefixime minimum inhibitory concentrations (MICs) ≥ 0.125 mg/L and ≥ 0.25 mg/L, respectively, were first determined in 259 strains isolated between 2013 and 2019 and then MICs of ertapenem were measured using the antimicrobial gradient epsilometer test (Etest). Genetic determinants of ESC resistance and multi-antigen sequence typing (NG-MAST) were also determined to analyze associations with ertapenem susceptibility.

**Results:** All isolates displayed ertapenem MICs between 0.006 mg/L-0.38 mg/L; the overall MIC_50_ and MIC_90_ were 0.032 mg/L and 0.125 mg/L. 44 (17.0%) isolates displayed ertapenem MICs of ≥ 0.125 mg/L; 10 (3.9%) had MICs ≥ 0.25 mg/L. The proportion of isolates with ertapenem MICs ≥ 0.125 mg/L increased from 4.0% in 2013, to 20.0% in 2019 (χ^2^= 24.144, P<0.001; Chi square test for linear trend). The *penA* mosaic allele was present in a significantly higher proportion of isolates with ertapenem MICs ≥ 0.125 mg/L compared to isolates with MICs ≤ 0.094 mg/L) (97.7% vs. 34.9%, respectively; χ^2^=58.158, P<0.001). ST5308 was the most prevalent NG-MAST type (8.5%); ST5308 was also significantly more common among isolates with ertapenem MICs ≥ 0.125 mg/L vs. isolates with MICs ≤ 0.094mg/L (22.7% and 5.6% respectively; χ^2^=13.815, P=0.001).

**Conclusions:** Ertapenem may be effective therapy for gonococcal isolates with decreased susceptibility or resistance to ESCs and isolates with identifiable genetic resistance determinants.

## INTRODUCTION

Gonorrhea is the second most common bacterial sexually transmitted infection and a major global public health problem. The World Health Organization (WHO) estimated that 87 million new cases occurred worldwide in adults aged 15-49, in 2016 (1). In China, the incidence of gonorrhea increased by 36.03% (7.05 to 9.59 cases per 100,000 population) from 2014 to 2018 (2). Treatment of gonorrhea is challenging because *N. gonorrhoeae* has developed resistance to most antimicrobials (AMR) that have been used for therapy, including sulfonamides, penicillins, tetracyclines, fluoroquinolones, early-generation and, rarely, extended-spectrum cephalosporins (ESCs) (3-7).

Currently, ceftriaxone monotherapy or dual therapy with ceftriaxone or cefixime plus azithromycin is recommended as first-line treatment of uncomplicated gonorrhea in most countries (8-10). Unfortunately, strains resistant to ESCs and azithromycin are emerging globally (9, 11) and treatment failures with currently-recommended dual therapies have been reported (12, 13). Thus, currently recommended treatments are unlikely to continue to be effective long-term; exploring novel or repurposed antimicrobials will be essential for control of gonorrhea (7).

Ertapenem is a parenteral carbapenem, effective against gram-negative bacteria that may, otherwise, be resistant to cephalosporins. Similar to other β-lactams, ertapenem inhibits cell wall synthesis by binding to and inhibiting penicillin-binding proteins (PBPs) (14). It is well tolerated, effective and has a safety profile comparable to that of ceftriaxone (15, 16). Ertapenem has been used successfully to treat *N. gonorrhoeae* with both high-level azithromycin and ceftriaxone resistance (13) and may be an effective treatment option for gonorrhea, particularly infections caused by strains resistant to extended spectrum cephalosporins (ESCs).

No specific genetic determinants of ertapenem resistance or carbapenemases, generally, have been identified in *N. gonorrhoeae*; however, there may be overlap with resistant mechanisms exhibited by other ESCs (17). Mechanisms of resistance against ESCs can result from amino acid changes caused by nucleotide mutations in: *penA* (encoding penicillin-binding protein 2, PBP2); *mtrR* (encoding the multiple transfer resistance repressor, MtrR); *penB* (encoding porin PorB) and *ponA* (encoding penicillin-binding protein 1, PBP1) in *N. gonorrhoeae* (3, 18-21).

The major aim of the present study was to examine *in vitro* activity of ertapenem, against *N. gonorrhoeae* isolates with decreased susceptibility (or resistance) to ESCs. We also identified ESC resistance determinants and their association with susceptibility of *N. gonorrhoeae* strains to ertapenem. Multiantigen sequence typing (NG-MAST) of *N. gonorrhoeae* isolates was performed to assess distribution according to ertapenem MICs and, potentially, to identify clonality of isolates with increased resistance.

## RESULTS

### Antimicrobial susceptibility

A total of 259 *N. gonorrhoeae* isolates with decreased susceptibility or resistance to ceftriaxone and/or cefixime were identified. The MIC ranges of ceftriaxone and cefixime for these isolates were 0.06-1 mg/L (MIC_50_, 0.125 mg/L and MIC_90_, 0.125 mg/L) and 0.06-≥4 mg/L (MIC_50_, 0.125 mg/L and MIC_90_, 0.5 mg/L), respectively. Among these isolates, 9 (3.5%) were fully resistant to ceftriaxone (MICs ≥0.5 mg/L) and cefixime (MICs ≥2 mg/L).

MICs of ertapenem against the 259 isolates ranged from 0.006 mg/L to 0.38 mg/L; MIC_50_ and MIC_90_ were 0.032 mg/L and 0.125 mg/L. For the 9 *N. gonorrhoeae* isolates fully resistant to ceftriaxone (MICs ≥ 0.5 mg/L) and cefixime (MICs ≥ 2 mg/L), the ertapenem MIC_50_, MIC_90_ and MIC range were 0.094, 0.19 and 0.023-0.19 mg/L, respectively. 44 (17.0%) isolates had ertapenem MICs ≥ 0.125 mg/L); 10 (3.9%) had MICs ≥ 0.25 mg/L, MICs that represent the WHO-recommended susceptibility breakpoints for ceftriaxone and cefixime, respectively. The ertapenem MIC_50_ and MIC_90_ increased from 0.023 mg/L and 0.047 mg/L in 2013 to 0.047 mg/L and 0.125 mg/L in 2019, respectively. The distributions of ertapenem MICs during 2013–2019 are shown in Figure 1. The proportion of isolates with ertapenem MICs ≥ 0.125 mg/L (the breakpoint against ceftriaxone) increased from 4.0% in 2013 to 20.0% in 2019, showing an overall upward trend during the study period (χ^2^= 24.144, P<0.001; Chi square test for linear trend), while the percent of isolates with MICs ≤ 0.012 mg/L declined in each successive year, sequentially (χ^2^= 23.634, P<0.001; Chi square test for linear trend).

**Figure 1.**
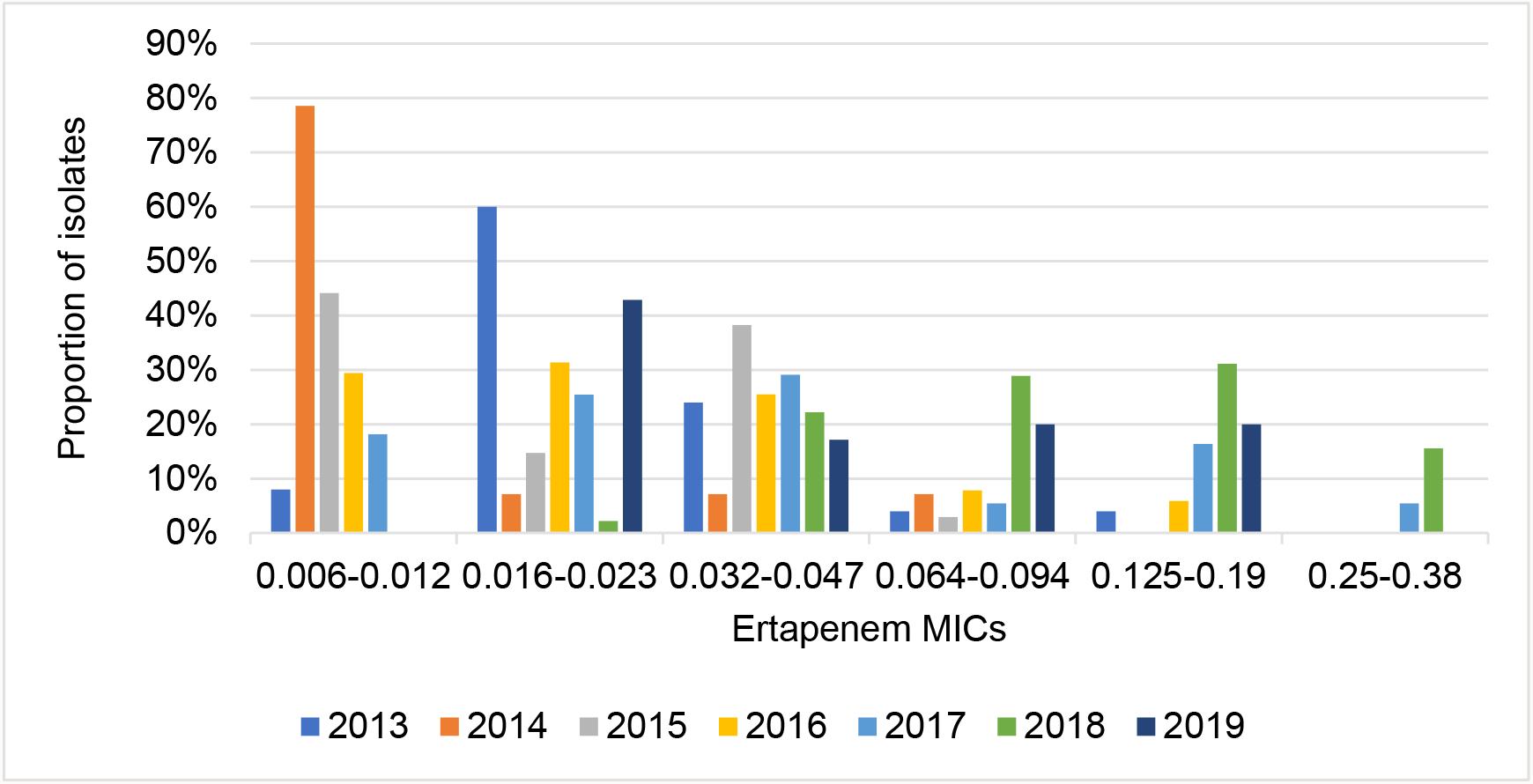
Distribution of ertapenem MICs for 259 *Neisseria gonorrhoeae* clinical isolates with decreased susceptibility (or resistance) to ESCs isolated in Nanjing, China, 2013–2019.

### Genetic resistance determinants (*penA, mtrR, penB* and *ponA*) of ESCs

A *penA* mosaic allele was present in 118 (45.6%) *N. gonorrhoeae* isolates with decreased susceptibility or resistance to ESCs; non-mosaic *penA* alleles with A501V/T mutations were present in 139 (53.7%); the remaining 2 isolates (0.8%) possessed a non-mosaic allele with an A517G mutation. Mutations in the promoter and/or coding regions of the *mtrR* gene were identified in 179 (69.1%) isolates. Amino acid substitutions at residue G120 of the *penB* gene were present in 5 (1.9%) isolates; G120/A121 double mutations were present in 253 (97.7%). An L421P mutation in the *ponA* gene was present in 256 (98.8%) isolates.

Ertapenem susceptibilities of isolates containing the *penA* mosaic allele, were lower compared to susceptibilities of isolates that lacked the mosaic allele. The MIC_50_, MIC_90_ and the MIC range of ertapenem in strains with the *penA* mosaic allele were 0.047 mg/L, 0.19 mg/L and 0.008-0.38 mg/L, respectively. Strains that lacked the mosaic allele had MIC_50_, MIC_90_ and an MIC range of ertapenem of 0.016 mg/L, 0.064 mg/L and 0.006–0.125 mg/L, respectively. The *penA* mosaic allele was more common among isolates with increased ertapenem MICs (≥ 0.125 mg/L) (WHO-recommended susceptibility breakpoint against ceftriaxone) vs. isolates with MICs ≤ 0.094 mg/L (97.7% vs. 34.9%, respectively; χ^2^=58.158, P<0.001; Table 1). All isolates with ertapenem MICs ≥ 0.25 mg/L (WHO-recommended susceptibility breakpoint against cefixime) possessed the *penA* mosaic allele. Conversely, the proportion of isolates with ertapenem MICs ≤ 0.094 mg/L that possessed A501V/T mutations, specifically, was higher than in isolates with MICs ≥ 0.125 mg/L (64.2% vs. 2.3%, respectively; χ^2^=56.307, P<0.001; Table 1). The two isolates with A517G mutations had ertapenem MICs of ≤ 0.094 mg/L (Table 1).

**Table 1.** *penA* and *mtrR* mutations in isolates either with MICs ≤0.094mg/L to ertapenem or MICs to ertapenem ≥0.125mg/L.

| | Number / proportion of isolates | | $\chi^2$ | P value <sup>a</sup> |
| --- | --- | --- | --- | --- |
| Resistance determinants | MIC≤0.094mg/L | MIC≥0.125mg/L |  |  |
|  | (N=215) | (N=44) |  |  |
| *penA* |  |  |  |  |
| Mosaic allele | 75 (34.9%) | 43 (97.7%) | 58.158 | <0.001 |
| A501V/T <sup>b</sup> | 138 (64.2%) | 1 (2.3%) | 56.307 | <0.001 |
| A517G <sup>b</sup> | 2(0.9%) | 0 | 0.413 | 1 |
| *mtrR* |  |  |  |  |
| A-deletion in promoter region <sup>c</sup> | 102 (47.4%) | 10 (22.7%) | 9.090 | 0.0026 |
| A-deletion <sup>c</sup>, A39T | 3 (1.4%) | 3 (6.8%) | 4.747 | 0.0632 |
| A-deletion <sup>c</sup>, G45D | 24 (11.2%) | 0 (0) | 5.413 | 0.0186 |
| A39T | 8 (3.7%) | 1 (2.3%) | 0.228 | 1.0000 |
| G45D | 27 (12.6%) | 1 (2.3%) | 4.007 | 0.0585 |
| WT | 51 (23.7%) | 29 (65.9%) | 30.453 | <0.001 |

<sup>a</sup> P<0.05 was considered significant in Chi-square ( $\chi^2$ ) or Fisher exact testing.
<sup>b</sup> Non-mosaic *penA* alleles.
<sup>c</sup> A (adenine) deletion in the 13-bp inverted-repeat sequence of the *mtrR* promoter.

*mtrR* mutations were present in 34.1% (15/44) of isolates with ertapenem MICs ≥ 0.125 mg/L and in 76.3% (164/215) of isolates with ertapenem MICs ≤0.094 mg/L (χ^2^= 30.453, P<0.001). A single A-deletion in the *mtrR* promoter was identified more often in isolates with ertapenem MICs ≤ 0.094 mg/L than in isolates with MICs ≥ 0.125 mg/L (χ^2^= 9.090, P=0.0026; Table 1). There were no significant differences in the rates of A39T or G45D *mtrR* mutations in the coding region accompanied (or not) by an A deletion in the promoter region. An exception was a G45D mutation accompanied by an A deletion in the promoter, which accounted for 11.2% (24/215) of isolates with ertapenem MICs ≤ 0.094 mg/L and no isolates with ertapenem MICs ≥ 0.125 mg/L (χ^2^= 5.413, P=0.0186; Table 1). All but two isolates with ertapenem MICs ≥ 0.25mg/L lacked the *mtrR* mutations; the two exceptions harbored a single A-deletion in the *mtrR* promoter or G45D mutation in the *mtrR* coding region.

### *N. gonorrhoeae* multiantigen sequence typing (NG-MAST)

The 259 *N. gonorrhoeae* isolates were assigned to 161 NG-MAST types, of which 68 have not been reported previously in the NG-MAST database (www.ng-mast.net). The most prevalent NG-MAST sequence type (ST) was ST5308 (n = 22; ertapenem MIC_50_, 0.094mg/L), followed by ST7554 (n=17; ertapenem MIC_50_, 0.032 mg/L), ST3356 (n = 7; ertapenem MIC_50_, 0.023 mg/L), ST270 (n = 7; ertapenem MIC_50_, 0.008 mg/L), and ST4539 (n = 7; ertapenem MIC_50_, 0.016 mg/L). Among all sequence types, ST5308 was predominant among isolates with MICs ≥ 0.125 mg/L to ertapenem (10/44 [22.7%]); these isolates also showed the highest ertapenem MIC_50_ (0.094mg/L). Furthermore, ST5308 was more common among isolates with MICs ≥ 0.125 mg/L to ertapenem vs. isolates with MICs ≤ 0.094 mg/L (22.7% and 5.6% respectively; χ^2^=13.815, P=0.001). All ST5308 isolates had a *penA* mosaic allele, G120K plus A121D substitutions in *penB* and L421P in *ponA* but no *mtrR* mutations.

## DISCUSSION

*Neisseria gonorrhoeae* is becoming increasingly resistant to currently used antimicrobial agents with the real prospect that untreatable gonorrhea may soon appear (9, 13). In the context of limited treatment options, alternative antimicrobials, new and repurposed, are needed urgently to ensure future successful treatments. Ertapenem is a member of the carbapenem family of antibiotics with a broad spectrum of activity. It is active against a variety of gram-positive and -negative bacteria, including *N. gonorrhoeae*. Ertapenem has been used successfully to treat *N. gonorrhoeae* that possessed combined high-level azithromycin and ceftriaxone resistance (13). In our study, the MIC_50_ of ertapenem (0.032 mg/L) was substantially lower than those observed for both ceftriaxone and cefixime (0.125 mg/L). The MIC_90_ of ertapenem (0.125 mg/L) was similar to the MIC_90_ observed for ceftriaxone (0.125 mg/L), but lower than the cefixime MIC_90_ (0.5 mg/L). Unemo et al (17) reported in 2012, that generally, ertapenem and ceftriaxone MIC_50_s and MIC_90_s were similar (0.032mg/L [both] and 0.64 mg/L [ertapenem] /0.125 mg/L [ceftriaxone], respectively) in 257 *N. gonorrhoeae* clinical isolates with highly diverse ceftriaxone MIC values referred to WHO Collaborating Centres for STIs. Similarly, ertapenem MIC ranges were lower than MIC ranges for ceftriaxone and cefixime in our study, also reported by Unemo et al (17). In our study, 83.0%/ 96.1% of isolates had ertapenem MICs below the ceftriaxone/cefixime breakpoints (0.125mg/L / 0.25 mg/L), similar to the study by Xu et al (22) that examined gonococcal isolates from eight provinces in China. In that study, 83.3% of 24 isolates with decreased susceptibility to ceftriaxone (MIC ≥ 0.25 mg/L) exhibited ertapenem MIC values <0.25 mg/L, the cefixime breakpoint. Unemo et al (17) reported that all strains had ertapenem MICs ≤ 0.125 mg/L, the ceftriaxone breakpoint. These results predict that ertapenem may be uniformly effective clinically in most instances because higher MICs are infrequent (our study and the study by Xu et al (22)) or absent altogether (17). Further support for clinical efficacy is derived from activity of ertapenem against two extensively drug resistant (XDR) *N. gonorrhoeae* strains, H041 and F89; both are highly resistant to cefixime (MIC range, 4-8 mg/L) and ceftriaxone (MIC range, 2-4 mg/L) (17). Ertapenem MICs were reported to be significantly lower (0.064 mg/L and 0.016 mg/L) for these two strains, respectively (17), corroborated in a separate study where F89 had an ertapenem MIC of 0.03 mg/L (23). In our study, ertapenem was also effective against the 9 *N. gonorrhoeae* isolates fully resistant to ceftriaxone (MICs ≥ 0.5 mg/L) and cefixime (MICs ≥ 2 mg/L). Nonetheless, these studies (17) (23) and another (24) have shown that ertapenem had no apparent *in vitro* advantage over ceftriaxone for *N. gonorrhoeae* isolates with lower ceftriaxone MICs.

Similar to other β-lactam antimicrobials, reduced activity of ertapenem against some bacteria is mediated by: mutations in porin that result in aberrant function (25); upregulation of efflux pumps (26) and production of carbapenemases (27). However, resistance of *N. gonorrhoeae* to ertapenem is not fully defined. For example, we found that, *penB* and *ponA* resistance determinants were present across most strains, perhaps without a meaningful effect on ertapenem susceptibility. We also found that the presence of a *penA* mosaic allele was strongly associated with increased MICs of ertapenem, similar to findings reported by Unemo et al (17). However, our result showed that *mtrR* mutations were present in a higher percentage of isolates with ertapenem MICs ≤ 0.094mg/L than in isolates with ertapenem MICs ≥ 0.125mg/L, different from another Chinese study, which showed that the *mtrR* promoter A-deletion was significantly associated with strains displaying an ertapenem MIC > 0.125 mg/L (28).

NG-MAST has been evaluated as a tool for predicting specific antimicrobial resistance phenotypes in *N. gonorrhoeae* isolates (29, 30). In our study, ST5308 was the most prevalent NG-MAST sequence type (ST) among the 259 isolates with decreased susceptibility or resistance to ESCs. In addition, ST5308 was the most highly represented ST in isolates with increased ertapenem MICs (≥ 0.125mg/L). ST5308 isolates, possessing a *penA* mosaic allele, have been reported in Hong Kong, and were associated with decreased susceptibility to oral ESCs (31). Between 2013 and 2017, ST5308 was the most common gonococcal type isolated in Guangdong, China (32).

In summary, *in vitro* susceptibility to ertapenem of *Neisseria gonorrhoeae* isolates with decreased susceptibility (or resistance) to ESCs, suggests potential for future use of ertapenem as treatment for antimicrobial resistant infections. However, the *penA* mosaic allele, commonly associated with ESC resistance, was also associated with increased MICs of ertapenem. Continued surveillance of antimicrobial susceptibility of ertapenem supplemented by sequence typing and NG-MAST classification are warranted.

## MATERIALS AND METHODS

### Bacterial strains

From January 2013 to December 2019, a total of 1321 *N. gonorrhoeae* strains were isolated from men with symptomatic urethritis (urethral discharge and/or dysuria) attending the STD clinic at the Institute of Dermatology, Chinese Academy of Medical Sciences, Nanjing, Jiangsu Province, China. Urethral exudates were collected with cotton swabs and immediately streaked on to modified Thayer-Martin (T-M) selective medium (Zhuhai DL Biotech Co. Ltd.) and incubated at 36°C in candle jars for 24–48 h. *N. gonorrhoeae* was identified by colonial morphology, Gram’s stain, and oxidase testing, which are sufficient to identify *N. gonorrhoeae* colonies isolated on selective medium, particularly for urethral samples from symptomatic men (33, 34). Gonococcal isolates were subcultured onto chocolate agar plates; pure colonies were swabbed, suspended in tryptone-based soy broth and frozen (−80°C) until used for antimicrobial testing.

### Antimicrobial susceptibility testing

Susceptibility testing for ceftriaxone and cefixime was performed by the agar dilution method according to Clinical and Laboratory Standards Institute (CLSI) guidelines (35). According to criteria for decreased susceptibility or resistance to ceftriaxone (MIC ≥ 0.125 mg/L) and cefixime (MIC ≥ 0.25 mg/L), defined by WHO (36), 259 strains were eligible for inclusion in this study. Ertapenem susceptibility among these isolates was determined by Etest (Liofilchem®, Italy) method, according to the manufacturer’s instructions (37). Strain WHO-P was used for quality control. No interpretative criteria have been provided by CLSI (or any other organization) for ertapenem susceptibility breakpoints against *N. gonorrhoeae*.

### Sequencing of resistance determinants (*penA, mtrR, penB* and *ponA*) and *N. gonorrhoeae* multiantigen sequence typing (NG-MAST)

Genomic DNA was prepared from individual gonococcal isolates using the Rapid Bacterial Genomic DNA Isolation Kit (DNA-EZ Reagents V All-DNA-Fast-Out, Sangon Biotech Co. Ltd, Shanghai). ESCs resistance determinants: *penA*; *mtrR*; *penB* and *ponA* were amplified by PCR using published primers (38) and DNA sequencing performed by Suzhou Genewiz Biotech Co. Ltd. The sequencing data was uploaded to the NG-STAR database (https://ngstar.canada.ca) to determine the ESCs resistance determinants.

Genetic characterization was performed by *N. gonorrhoeae* multiantigen sequence typing (NG-MAST), which assigns sequence types (STs) based on a combination of two variable genes: *porB* and *tbpB* (39); allele numbers and sequence types (STs) were then assigned (http://www.ng-mast.net).

### Data Analysis

Chi-square (χ^2^) testing for linear trends was used to assess changes in ertapenem MICs during the study period. Chi-square or Fisher exact testing was used to determine the associations between ertapenem susceptibility and gonococcal genetic resistance determinants or *N. gonorrhoeae* multi-antigen sequence types. SPSS version 26.0 was used for statistical analysis and P values <0.05 considered significant.

## ACKNOWLEDGEMENTS

This work was supported by grants from the Chinese Academy of Medical Sciences Initiative for Innovative Medicine (2016-I 2M-3-021) and the U.S. National Institutes of Health (AI084048 and AI116969).

## REFERENCES

1. World Health Organization. 2018. Report on global sexually transmitted infection surveillance -2018. World Health Organization, Geneva, Switzerland.

2. Yue, X., X. Gong, J. Li, Y. Wang, and H. Gu. 2019. Gonorrhea in China, 2018. International Journal of Dermatology and Venereology 2:65–69.

3. Ohnishi, M., D. Golparian, K. Shimuta, T. Saika, S. Hoshina, K. Iwasaku, S. Nakayama, J. Kitawaki, and M. Unemo. 2011. Is Neisseria gonorrhoeae initiating a future era of untreatable gonorrhea?: detailed characterization of the first strain with high-level resistance to ceftriaxone. Antimicrob Agents Chemother 55:3538–45.

4. Unemo, M., D. Golparian, R. Nicholas, M. Ohnishi, A. Gallay, and P. Sednaoui. 2012. High-level cefixime-and ceftriaxone-resistant Neisseria gonorrhoeae in France: novel penA mosaic allele in a successful international clone causes treatment failure. Antimicrob Agents Chemother 56:1273–80.

5. Camara, J., J. Serra, J. Ayats, T. Bastida, D. Carnicer-Pont, A. Andreu, and C. Ardanuy. 2012. Molecular characterization of two high-level ceftriaxone-resistant Neisseria gonorrhoeae isolates detected in Catalonia, Spain. J Antimicrob Chemother 67:1858–60.

6. Unemo, M., C. S. Bradshaw, J. S. Hocking, H. de Vries, S. C. Francis, D. Mabey, J. M. Marrazzo, G. Sonder, J. R. Schwebke, E. Hoornenborg, R. W. Peeling, S. S. Philip, N. Low, and C. K. Fairley. 2017. Sexually transmitted infections: challenges ahead. Lancet Infect Dis 17:e235–e279.

7. Wi, T., M. M. Lahra, F. Ndowa, M. Bala, J. R. Dillon, P. Ramon-Pardo, S. R. Eremin, G. Bolan, and M. Unemo. 2017. Antimicrobial resistance in Neisseria gonorrhoeae: Global surveillance and a call for international collaborative action. PLoS Med 14:e1002344.

8. World Health Organization. 2016. WHO guidelines for the treatment of Neisseria gonorrhoeae. World Health Organization, Geneva, Switzerland.

9. Unemo, M., and W. M. Shafer. 2014. Antimicrobial resistance in Neisseria gonorrhoeae in the 21st century: past, evolution, and future. Clin Microbiol Rev 27:587–613.

10. St, C. S., L. Barbee, K. A. Workowski, L. H. Bachmann, C. Pham, K. Schlanger, E. Torrone, H. Weinstock, E. N. Kersh, and P. Thorpe. 2020. Update to CDC’s Treatment Guidelines for Gonococcal Infection, 2020. MMWR Morb Mortal Wkly Rep 69:1911–1916.

11. Yan, J., J. Xue, Y. Chen, S. Chen, Q. Wang, C. Zhang, S. Wu, H. Lv, Y. Yu, and S. van der Veen. 2019. Increasing prevalence of Neisseria gonorrhoeae with decreased susceptibility to ceftriaxone and resistance to azithromycin in Hangzhou, China (2015-17). J Antimicrob Chemother 74:29–37.

12. Fifer, H., U. Natarajan, L. Jones, S. Alexander, G. Hughes, D. Golparian, and M. Unemo. 2016. Failure of Dual Antimicrobial Therapy in Treatment of Gonorrhea. N Engl J Med 374:2504–6.

13. Eyre, D. W., N. D. Sanderson, E. Lord, N. Regisford-Reimmer, K. Chau, L. Barker, M. Morgan, R. Newnham, D. Golparian, M. Unemo, D. W. Crook, T. E. Peto, G. Hughes, M. J. Cole, H. Fifer, A. Edwards, and M. I. Andersson. 2018. Gonorrhoea treatment failure caused by a Neisseria gonorrhoeae strain with combined ceftriaxone and high-level azithromycin resistance, England, February 2018. Euro Surveill 23.

14. Congeni, B. L. 2010. Ertapenem. Expert Opin Pharmacother 11:669–72.

15. Arguedas, A., J. Cespedes, F. A. Botet, J. Blumer, R. Yogev, R. Gesser, J. Wang, J. West, T. Snyder, and W. Wimmer. 2009. Safety and tolerability of ertapenem versus ceftriaxone in a double-blind study performed in children with complicated urinary tract infection, community-acquired pneumonia or skin and soft-tissue infection. Int J Antimicrob Agents 33:163–7.

16. Burkhardt, O., H. Derendorf, and T. Welte. 2007. Ertapenem: the new carbapenem 5 years after first FDA licensing for clinical practice. Expert Opin Pharmacother 8:237–56.

17. Unemo, M., D. Golparian, A. Limnios, D. Whiley, M. Ohnishi, M. M. Lahra, and J. W. Tapsall. 2012. In vitro activity of ertapenem versus ceftriaxone against Neisseria gonorrhoeae isolates with highly diverse ceftriaxone MIC values and effects of ceftriaxone resistance determinants: ertapenem for treatment of gonorrhea? Antimicrob Agents Chemother 56:3603–9.

18. Golparian, D., B. Hellmark, H. Fredlund, and M. Unemo. 2010. Emergence, spread and characteristics of Neisseria gonorrhoeae isolates with in vitro decreased susceptibility and resistance to extended-spectrum cephalosporins in Sweden. Sex Transm Infect 86:454–60.

19. Ison, C. A., J. Hussey, K. N. Sankar, J. Evans, and S. Alexander. 2011. Gonorrhoea treatment failures to cefixime and azithromycin in England, 2010. Euro Surveill 16.

20. Lindberg, R., H. Fredlund, R. Nicholas, and M. Unemo. 2007. Neisseria gonorrhoeae isolates with reduced susceptibility to cefixime and ceftriaxone: association with genetic polymorphisms in penA, mtrR, porB1b, and ponA. Antimicrob Agents Chemother 51:2117–22.

21. Zhao, S., M. Duncan, J. Tomberg, C. Davies, M. Unemo, and R. A. Nicholas. 2009. Genetics of chromosomally mediated intermediate resistance to ceftriaxone and cefixime in Neisseria gonorrhoeae. Antimicrob Agents Chemother 53:3744–51.

22. Xu, W. Q., X. L. Zheng, J. W. Liu, Q. Zhou, X. Y. Zhu, J. Zhang, Y. Han, K. Chen, S. C. Chen, X. S. Chen, and Y. P. Yin. 2021. Antimicrobial Susceptibility of Ertapenem in Neisseria gonorrhoeae Isolates Collected Within the China Gonococcal Resistance Surveillance Programme (China-GRSP) 2018. Infect Drug Resist 14:4183–4189.

23. Quaye, N., M. J. Cole, and C. A. Ison. 2014. Evaluation of the activity of ertapenem against gonococcal isolates exhibiting a range of susceptibilities to cefixime. J Antimicrob Chemother 69:1568–71.

24. Livermore, D. M., S. Alexander, B. Marsden, D. James, M. Warner, E. Rudd, and K. Fenton. 2004. Activity of ertapenem against Neisseria gonorrhoeae. J Antimicrob Chemother 54:280–1.

25. Jacoby, G. A., D. M. Mills, and N. Chow. 2004. Role of beta-lactamases and porins in resistance to ertapenem and other beta-lactams in Klebsiella pneumoniae. Antimicrob Agents Chemother 48:3203–6.

26. Szabo, D., F. Silveira, A. M. Hujer, R. A. Bonomo, K. M. Hujer, J. W. Marsh, C. R. Bethel, Y. Doi, K. Deeley, and D. L. Paterson. 2006. Outer membrane protein changes and efflux pump expression together may confer resistance to ertapenem in Enterobacter cloacae. Antimicrob Agents Chemother 50:2833–5.

27. McGettigan, S. E., K. Andreacchio, and P. H. Edelstein. 2009. Specificity of ertapenem susceptibility screening for detection of Klebsiella pneumoniae carbapenemases. J Clin Microbiol 47:785–6.

28. Yang, F., J. Yan, J. Zhang, and S. van der Veen. 2020. Evaluation of alternative antibiotics for susceptibility of gonococcal isolates from China. Int J Antimicrob Agents 55:105846.

29. Palmer, H. M., H. Young, C. Graham, and J. Dave. 2008. Prediction of antibiotic resistance using Neisseria gonorrhoeae multi-antigen sequence typing. Sex Transm Infect 84:280–4.

30. Thakur, S. D., P. N. Levett, G. B. Horsman, and J. A. Dillon. 2014. Molecular epidemiology of Neisseria gonorrhoeae isolates from Saskatchewan, Canada: utility of NG-MAST in predicting antimicrobial susceptibility regionally. Sex Transm Infect 90:297–302.

31. Lo, J. Y., K. M. Ho, and A. C. Lo. 2012. Surveillance of gonococcal antimicrobial susceptibility resulting in early detection of emerging resistance. J Antimicrob Chemother 67:1422–6.

32. Qin, X., Y. Zhao, W. Chen, X. Wu, S. Tang, G. Li, Y. Yuqi, W. Cao, X. Liu, J. Huang, J. Yang, W. Chen, W. Tang, and H. Zheng. 2019. Changing antimicrobial susceptibility and molecular characterisation of Neisseria gonorrhoeae isolates in Guangdong, China: in a background of rapidly rising epidemic. Int J Antimicrob Agents 54:757–765.

33. Ison, C. A. 1990. Laboratory methods in genitourinary medicine. Methods of diagnosing gonorrhoea. Genitourin Med 66:453–9.

34. Unemo M and Ison C. 2013. Gonorrhoea, p 21–54. In Laboratory diagnosis of sexually transmitted infections, including human immunodeficiency virus. World Health Organization (WHO), Geneva, Switzerland.

35. Clinical and Laboratory Standards Institute. 2012. Performance standards for antimicrobial susceptibility testing; 22nd informational supplement. CLSI document M100-S22. Clinical and Laboratory Standards Institute, Wayne, PA.

36. World Health Organization (WHO), Department of Reproductive Health and Research. 2012. Global action plan to control the spread and impact of antimicrobial resistance in Neisseria gonorrhoeae, p 1–36. WHO, Geneva, Switzerland.

37. AB Biodisk. Etest application sheet. M0000503-MH0184 AB Biodisk, Solna, Sweden 2007.Available from: http://www.ilexmedical.com/files/EtestApplicationSheets.pdf.

38. Demczuk, W., S. Sidhu, M. Unemo, D. M. Whiley, V. G. Allen, J. R. Dillon, M. Cole, C. Seah, E. Trembizki, D. L. Trees, E. N. Kersh, A. J. Abrams, H. de Vries, A. P. van Dam, I. Medina, A. Bharat, M. R. Mulvey, G. Van Domselaar, and I. Martin. 2017. Neisseria gonorrhoeae Sequence Typing for Antimicrobial Resistance, a Novel Antimicrobial Resistance Multilocus Typing Scheme for Tracking Global Dissemination of N. gonorrhoeae Strains. J Clin Microbiol 55:1454–1468.

39. Martin, I. M., C. A. Ison, D. M. Aanensen, K. A. Fenton, and B. G. Spratt. 2004. Rapid sequence-based identification of gonococcal transmission clusters in a large metropolitan area. J Infect Dis 189:1497–505.

